# Genomic surveillance of SARS-CoV-2 reveals community transmission of a major lineage during the early pandemic phase in Brazil

**DOI:** 10.1101/2020.06.17.158006

**Authors:** Paola Cristina Resende, Edson Delatorre, Tiago Gräf, Daiana Mir, Fernando do Couto Motta, Luciana Reis Appolinario, Anna Carolina Dias da Paixão, Maria Ogrzewalska, Braulia Caetano, Mirleide Cordeiro dos Santos, Jessylene de Almeida Ferreira, Edivaldo Costa Santos Junior, Sandro Patroca da Silva, Sandra Bianchini Fernandes, Lucas A Vianna, Larissa da Costa Souza, Jean F G Ferro, Vanessa B Nardy, Júlio Croda, Wanderson K Oliveira, André Abreu, Gonzalo Bello, Marilda M Siqueira

## Abstract

Despite all efforts to control the COVID-19 spread, the SARS-CoV-2 reached South America within three months after its first detection in China, and Brazil became one of the hotspots of COVID-19 in the world. Several SARS-CoV-2 lineages have been identified and some local clusters have been described in this early pandemic phase in Western countries. Here we investigated the genetic diversity of SARS-CoV-2 during the early phase (late February to late April) of the epidemic in Brazil. Phylogenetic analyses revealed multiple introductions of SARS-CoV-2 in Brazil and the community transmission of a major B.1.1 lineage defined by two amino acid substitutions in the Nucleocapsid and ORF6. This SARS-CoV-2 Brazilian lineage was probably established during February 2020 and rapidly spread through the country, reaching different Brazilian regions by the middle of March 2020. Our study also supports occasional exportations of this Brazilian B.1.1 lineage to neighboring South American countries and to more distant countries before the implementation of international air travels restrictions in Brazil.

## Introduction

COVID-19, the disease caused by Severe Acute Respiratory Syndrome Coronavirus-2 (SARS-CoV-2), is leading to high rates of acute respiratory syndrome, hospitalization, and death ^1,2^. Brazil, the second most hit country in the world so far, has reported 923.189 cases and 45.241 deaths (last update 17^th^ June 2020) ^3^. The first positive case of SARS-CoV-2 infection in Brazil was reported on 26^th^ February 2020 in an individual traveling from Europe to Sao Paulo metropolitan region ^4^, and during the following two weeks, the virus was detected in all country regions ^5^

The rapid worldwide genomic surveillance of SARS-CoV-2, mainly shared via the GISAID (https://www.gisaid.org/) databank that provides public access to genomic sequence and patient’s metadata, is being crucial for managing this healthcare emergency enabling the tracking of viral transmission patterns as the epidemic progresses. The SARS-CoV-2 has diversified in several phylogenetic lineages while it spread geographically across the world ^6–8^. A SARS-CoV-2 lineage previously designated as “G” or “B.1” clade, was initially identified as the most common variant in Europe and is currently one of the predominant viral lineages in North America ^6–8^. Inspection of SARS-CoV-2 genomic sequences from South America available on GISAID revealed that the clade B.1 is also the most prevalent (82%) SARS-CoV-2 variant circulating in South America (**Fig. 1A**).

**Figure 1.**
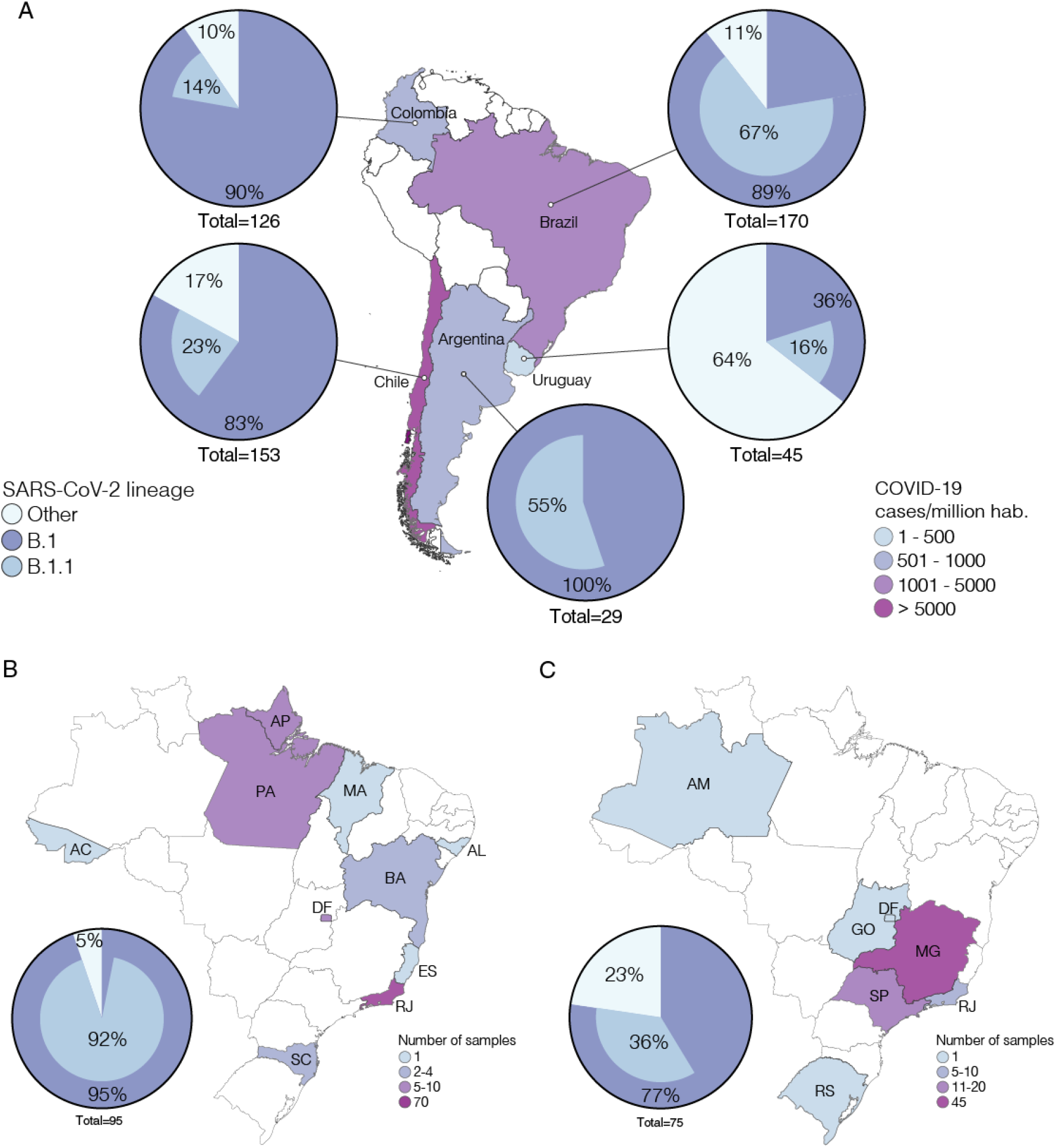
Prevalence of SARS-CoV-2 clades B.1 and B.1.1 in Brazil and other South American countries. A) Map showing the prevalence of SARS-CoV-2 clades B.1 and B.1.1 across different South American countries with more than five viral genomes available in the GISAID (https://www.gisaid.org/) database as of 4^th^ June. Countries were colored according to the incidence of COVID-19. B and C) Brazilian maps showing the prevalence of SARS-CoV-2 clades B.1 and B.1.1 across different states considering the viral sequences generated in this study (B) or those generated by others and deposited in the GISAID database. Brazilian states are colored according to the number of SARS-CoV-2 available. Colors in pie charts correspond to the viral lineage.

Genomic epidemiology has been a useful tool to track the community transmission of SARS-CoV-2 in different geographical settings. Previous studies revealed that SARS-CoV-2 epidemics in Australia ^9,10^, Belgium ^11^, Denmark ^12^, France ^13^, Iceland ^14^, Israel ^15^, Netherlands ^16^ Spain ^17^ and the United States (US) ^18–20^, resulted from multiple independent introductions, followed by community dissemination of some viral strains that resulted in the emergence of national (or local) transmission clusters. Early phylogenetic analyses of SARS-CoV-2 complete genomes from the Brazilian states of Minas Gerais ^21^, Sao Paulo ^22^ and Amazonas ^23^, revealed multiple independent viral importations and limited local spread during the initial stage of the SARS-CoV-2 epidemic in Brazil. Even so, the SARS-CoV-2 genomes analyzed in those previous studies were mostly recovered from individuals returning from international travel, and thus might not have recovered the genetic diversity of SARS-CoV-2 strains linked to community transmission in Brazil.

To investigate the SARS-CoV-2 strains circulating in Brazil, we recovered 95 whole-genomes collected from 10 different Brazilian states during the first two months of the COVID-19 epidemic. New SARS-CoV-2 Brazilian viral sequences were combined with other Brazilian and global reference sequences available in GISAID and subjected to maximum likelihood (ML) and Bayesian coalescent analyses.

## Methods

### Sampling and ethical aspects

Nasopharyngeal-throat combined swabs were collected from clinically ill individuals between the first and the eleventh day after their first symptoms, or from asymptomatic individuals suspicious of SARS-CoV-2 infection. Samples were conserved in the viral transport medium at 4°C to 8°C up to processing. This study was approved by the FIOCRUZ-IOC Ethics Committee (68118417.6.0000.5248) and the Brazilian Ministry of Health SISGEN (A1767C3).

### Nucleic acid isolation and RT–qPCR

The Viral RNA was extracted manually from 140 μl of clinical samples using QIAamp Viral RNA Mini kit (QIAGEN, Hilden, Germany) or automatedly using the 300 μl of sample and Perkin-Elmer Chemagic machine/chemistry, according to the manufacturer’s instructions. SARS-CoV-2 positive cases were confirmed by real-time RT–PCR assays using the SARS-COV-2 Molecular E/RP Kit (Biomanguinhos, Rio de Janeiro, Brazil) based on the protocol previously designed by Corman et al (2020) ^24^. Amplifications were conducted in ABI7500 platform using the following conditions: reverse transcription (50°\C, 15 min), reverse transcriptase inactivation and DNA polymerase activation (95 °C, 2 min), followed by 45 cycles of DNA denaturation (95 °C, 20 s) and annealing–extension (58 °C, 30 s). The fluorescence data was collected in the annealing-extension step and all samples with sigmoid curves crossing the threshold line up to cycle 40 were named positive. Negative and positive controls were included in each extraction and real time RT– PCR batch.

### SARS-CoV-2 whole-genome amplification and sequencing

Total RNA from positive samples presenting Ct values up to 30,0 for gene E was reverse transcribed using SuperScript™ IV First Strand Synthesis System (Invitrogen). Two multiplex PCR reactions using the primer scheme previously described ^25^ (Pool A = nine amplicons and Pool B = eight amplicons), were performed using the Q5® High-Fidelity DNA Polymerase (NEB). Amplicons were purified using Agencourt AMPure XP beads (Beckman Coulter™) and the DNA quantified with Qubit 4 Fluorometer (Invitrogen) using the Qubit dsDNA HS Assay Kit (Invitrogen) and sequenced using Illumina MiSeq or NextSeq (San Diego, CA, USA) and Nanopore (Oxford, UK) platforms. Illumina short reads DNA libraries were generated from the pooled amplicons using Nextera XT DNA Sample Preparation Kit (Illumina, San Diego, CA, USA) according to the manufacturer specifications. The size distribution of these libraries was evaluated using a 2100 Bioanalyzer (Agilent, Santa Clara, USA) and the samples were pair-end sequenced (Micro V2, 300 cycles) on a MiSeq equipment (Illumina, San Diego, USA) in around 18 hours. The Nanopore library protocol is optimized for long reads (2 kb amplicons)^25^. Library preparation was conducted using Ligation Sequencing 1D (SQK-LSK109 Oxford Nanopore Technologies (ONT) and Native Barcoding kit 1 to 24 (ONT), according to the manufacturer’s instructions. After end repair using the NEBNext® Ultra™ II End Repair/dA-Tailing Module (New England Biolabs, NEB) the native barcodes were attached using a NEBNext® Ultra™ II Ligation Module (NEB). Up to 23 samples were pooled for sequencing in the same flow cell (FLOMIN106 flow cell R9.4.1), and a negative mock sample was loaded in each run for validation. The sequencing was performed for 12 hours using the high accuracy base calling in the MinKNOW software, however, the run was monitored by RAMPART (https://github.com/articnetwork/rampart) allowing us stop the assay after 2 hours, when ≥ 20× depth for all amplicons was achieved.

### Data analysis to recover the SARS-CoV-2 whole-genome consensus sequences

Demultiplexed fastq files generated by Illumina sequencing were used as the input for the analysis. Reads were trimmed based on quality scores with a cutoff of q30, in order to remove low quality regions and adapter sequences. The reads were mapped to Wuhan Strain MN908947. Duplicate reads were removed from the alignment and the consensus sequence called at a threshold of 10×. The entire workflow was carried out in CLC Genomics Workbench software version 20.0. For the Oxford Nanopore sequencing data, the high accuracy base called fastq files were used as an input for analysis. The pipeline used was an adaptation of the artic-ncov2019 medaka workflow (https://artic.network/ncov-2019/ncov2019-bioinformatics-sop.html). We used an earlier version of the workflow which used Porechop to demultiplex the reads. The mapping to the Wuhan reference sequence (MN908947) was done using Minimap2 with Medaka used for error correction. This was all carried out within the artic-ncov2019-medaka conda environment (https://github.com/artic-network/artic-ncov2019).

### SARS-CoV-2 genotyping

New Brazilian genome sequences of SARS-CoV-2 were assigned to viral lineages according to the nomenclature proposed by Rambaut *et al ^7^*, using the pangolin web application (https://pangolin.cog-uk.io). A matrix with the count of each possible character at each position of the alignment of the B.1.1 sequences available in GISAID as of June 6, was computed using the R package SeqinR ^26^.

### Maximum Likelihood phylogenetic analyses

SARS-CoV-2 B.1.1 complete genome sequences (> 29 Kilobases) with appropriate metadata were retrieved from GISAID (https://www.gisaid.org/) as of 4^th^ June. After excluding low quality genomes (> 10% of ambiguous positions), we obtained a final dataset of 7,674 sequences. Because most sequences recovered (75%) were from the United Kingdom (UK), we generate a “non-redundant” global balanced dataset by removing very closely related sequences (genetic similarity 99.99%) from the UK. To achieve this aim, sequences from the UK were grouped by similarity with the CD-HIT program ^27^ and one sequence per cluster was selected. With this sampling procedure, we obtained a balanced global reference B.1.1 dataset containing 3,764 sequences that were aligned with the new B.1.1 Brazilian sequences generated in this study using MAFFT v7.467 ^28^ and then subjected to maximum-likelihood (ML) phylogenetic analyses. The ML phylogenetic tree was inferred using IQTREE v1.6.12 ^29^, under the GTR+F+I+G4 nucleotide substitution model as selected by the ModelFinder application ^30^ and the branch support was assessed by the approximate likelihood-ratio test based on a Shimodaira–Hasegawa-like procedure (SH-aLRT) with 1,000 replicates. The ML tree was visualized using the FigTree v1.4 (http://tree.bio.ed.ac.uk/software/figtree/).

### Analysis of temporal signal and phylogeographic structure

A ML tree of the B.1.1.EU/BR and B.1.1.BR dataset was inferred as explained above and the temporal signal was assessed by performing a regression analysis of the root-to-tip divergence against sampling time using TempEst ^31^. The degree of phylogeographic structure was then investigated using the BaTS program ^32^ which estimates phylogeny-trait associations in a posterior sampling of Bayesian trees. Bayesian trees were generated with BEAST package ^33^ as explained below, but without incorporation of a phylogeographic model. Phylogenetic clustering by sampling location in the posterior sampling of trees was then assessed by calculating different metrics including the Association Index (AI), the Parsimony Score (PS) and the Maximum Clade (MC) and compared to a null hypothesis generated by tip randomization. Results were considered significant for *P* < 0.01.

### Bayesian phylogeographic analyses

The age of the most recent common ancestor (*T*MRCA) and the spatial diffusion pattern of the B.1.1.EU/BR and B.1.1.BR lineages were jointly estimated using a Bayesian Markov Chain Monte Carlo (MCMC) approach implemented in BEAST 1.10 ^33^, using the BEAGLE library v3 ^34^ to improve computational time. Time-scaled Bayesian trees were estimated by using a strict molecular clock model with a fixed substitution rate (8 × 10^−4^ substitutions/site/year) based on previous estimates, the HKY+I+G nucleotide substitution model, and the Bayesian skyline coalescent prior ^*35*^. Viral migrations across time were reconstructed using a reversible discrete phylogeographic model ^36^ with a CTMC rate reference prior ^37^. Two MCMC chains were run for 100 million generations and then combined to ensure stationarity and good mixing. Stationarity (constant mean and variance of trace plots) and good mixing (Effective Sample Size >200) for all parameter estimates were assessed using TRACER v1.7 ^38^. The maximum clade credibility (MCC) tree was summarized with TreeAnnotator v1.10 and visualized using the FigTree v1.4 program.

## Results

In this study, we analyzed 95 viral whole-genomes (>99% coverage) obtained from individuals with confirmed SARS-CoV-2 infection, who underwent testing and genomic sequencing at the Laboratory of Respiratory Viruses and Measles, Oswaldo Cruz Institute (IOC)-FIOCRUZ, in Rio de Janeiro, and the Evandro Chagas Institute, in Para, Brazil ^25^. Samples were collected between 29^th^ February and 28^th^ April 2020 from individuals that reside in 10 different Brazilian states from the Southeastern (Rio de Janeiro and Espirito Santo), Central-Western (Distrito Federal), Northern (Acre, Amapa and Para), Northeastern (Alagoas, Bahia and Maranhao) and Southern (Santa Catarina) regions. (**Supplementary Table 1)**. The median age of patients with COVID-19 illness was 42-year-old (range 0 to 85 years) and 54 (57%) were female. Seven individuals reported international travel or contact with travelling people. Six different SARS-CoV-2 lineages (A.2, B.1, B.1.1, B.2.1, B.2.2 and B.6) were detected in our sample (**Supplementary Table 1),** according to the nomenclature proposed by Rambaut *et al ^7^*. Most Brazilian SARS-CoV-2 sequences here obtained were classified as clade B.1 (95%, n = 90), and particularly within the sub-clade B.1.1 (92%, n = 87) (**Fig. 1B**). The prevalence of the sub-clade B.1.1 in our sample (92%) was much higher than that observed in other Brazilian sequences available in GISAID (36%) (**Fig. 1C**). The clade B.1.1 was the only lineage detected in the 18 individuals with no history of recent international travel, while four different lineages (B.1, B.1.1, B2.1 and B.6) were detected among the six individuals with recent history of international travel (imported cases) and their contacts. (**Supplementary Table 1**).

To investigate whether the observed high prevalence of the lineage B.1.1 in Brazil resulted from one or multiple independent viral introductions into the country, we performed a ML phylogenetic analysis of the 87 B.1.1 Brazilian sequences identified in this study, together with 3,764 SARS-CoV-2 complete genome sequences available in GISAID as of 4^th^ June representing the current global diversity of the B.1.1 clade. Brazilian isolates were distributed throughout the phylogenetic tree, consistent with the hypothesis of multiple independent introductions (**Fig. 2**). A significant proportion of Brazilian B.1.1 sequences (65%, *n* = 74/114), however, branched in a monophyletic cluster (SH-*aLRT* = 74%) here designated as B.1.1.BR, that comprises sequences from Brazil, other South American countries (Argentina, Chile and Uruguay), North America (Canada and USA), Australia and England. The lineage B.1.1.BR is nested within a highly supported (SH-*aLRT* = 87%) clade, here referred as B.1.1.EU/BR, containing basal sequences from Western Europe and Brazil, (**Fig. 2**). We also detected two other well-supported (SH-*aLRT* 80%) monophyletic clades of small size (*n* = 2-11) mostly composed by Brazilian sequences (**Supplementary Fig. 1**).

**Figure 2.**
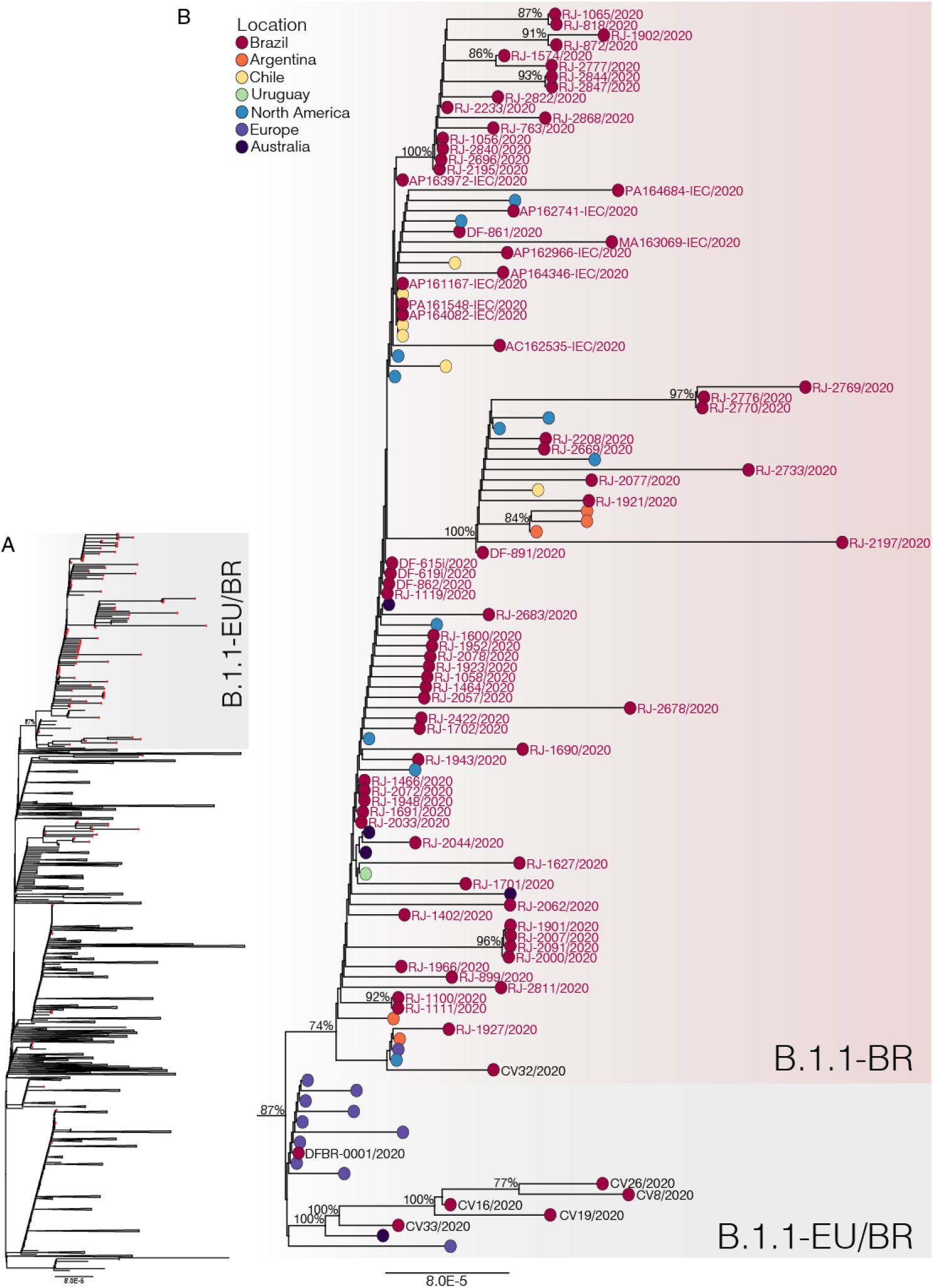
Phylogenetic relationships of SARS-CoV-2 B.1.1 Brazilian and global strains. A) ML phylogenetic tree of 87 B.1.1 Brazilian genomes obtained in this survey (red circles) along with 3,764 B.1.1 worldwide reference sequences from GISAID database. B) Zoomed view of the clusters B.1.1.EU/BR and B.1.1.BR. Names of Brazilian SARS-CoV-2 genomes generated in this study (red tips) or deposited in GISAID databank (black tips) are shown. Tip circles are colored according to the sampling location. Only node supports (aLRT) above 70% are shown. Shaded boxes highlight the position of clusters B.1.1.EU/BR and B.1.1.BR. Tree was rooted on midpoint and branch lengths are drawn to scale with the bars at the bottom indicating nucleotide substitutions per site.

In addition to sharing the three nucleotide mutations (G28881A, G28882A, G28883C) characteristic of the clade B.1.1, sequences of clusters B.1.1.EU/BR and B.1.1.BR harbor a non-synonymous mutation T29148C at the Nucleocapsid protein (I292T); and another non-synonymous mutation T27299C at the ORF6 (I33T) was shared only by sequences of the lineage B.1.1.BR. Mutations T29148C or T27299C were not detected in the other 7,551 B.1.1 genomes available in GISAID, supporting the hypothesis that they are synapomorphic traits of the B.1.1.EU/BR and B.1.1.BR clades, respectively. The clades B.1.1.EU/BR and B.1.1.BR were detected in different countries around the world, but the overall estimated prevalence of these clades in Brazil (6% and 44%, respectively) is much higher than that estimated in Europe, North America or Australia (**Supplementary Table 2**). Such difference could not be explained by sampling bias as those regions comprise the most densely sampled countries worldwide and is suggestive of local dissemination of those clades in Brazil. Consistent with this hypothesis, none of the individuals infected with clade B.1.1.BR from our cohort or with clade B.1.1.EU/BR from a previous cohort ^21^ reported international travel. The clades B.1.1.EU/BR and B.1.1.BR were not homogeneously distributed across Brazilian states (**Supplementary Table 3**). The clade B.1.1.EU/BR was highly prevalent in Minas Gerais and also detected in the Federal District, while the clade B.1.1.BR was predominant in Rio de Janeiro and also identified in some samples from the Northern, Central-Western and Northeastern Brazilian regions. Notably, none of these lineages were detected in the most populated state of Sao Paulo.

Finally, we conducted a Bayesian phylogeographic analysis to reconstruct the spatiotemporal dissemination dynamics of the B.1.1.EU/BR and B.1.1.BR lineages. Linear regression of root-to-tip genetic distance against sampling date revealed a weak temporal structure in our dataset (R^2^ = 0.19) (**Supplementary Fig. 2**). Despite the low genetic diversity, analyses of geographic structure rejected the null hypothesis of a panmixed population (**Supplementary Table 4**), supporting that geographic subdivision of the B.1.1.EU/BR and B.1.1.BR sequences was greater than expected by chance. The time-scaled Bayesian tree was then reconstructed using a strict molecular clock model with a fixed substitution rate (8 × 10^−4^ substitutions/site/year). Bayesian reconstructions traced the origin of the B.1.1.EU/BR lineage most probably to Europe (*Posterior state probability* [*PSP*] = 0.64) at 2^nd^ February (95% High Posterior Density [HPD]: 7^th^ January – 20^th^ February) and its dissemination to Brazil at 19^th^ February (95% HPD: 4^th^ February – 28^th^ February) (**Fig. 3A**). The origin of the B.1.1.BR lineage was traced with high probability to Brazil (*PSP* = 0.95) at 22^th^ February (95% HPD: 10^th^ February – 28^th^ February). From Brazil, the B.1.1.BR lineage probably disseminated to neighboring South American countries (Argentina, Chile and Uruguay) and to more distant regions (Australia, USA and UK).

**Figure 3.**
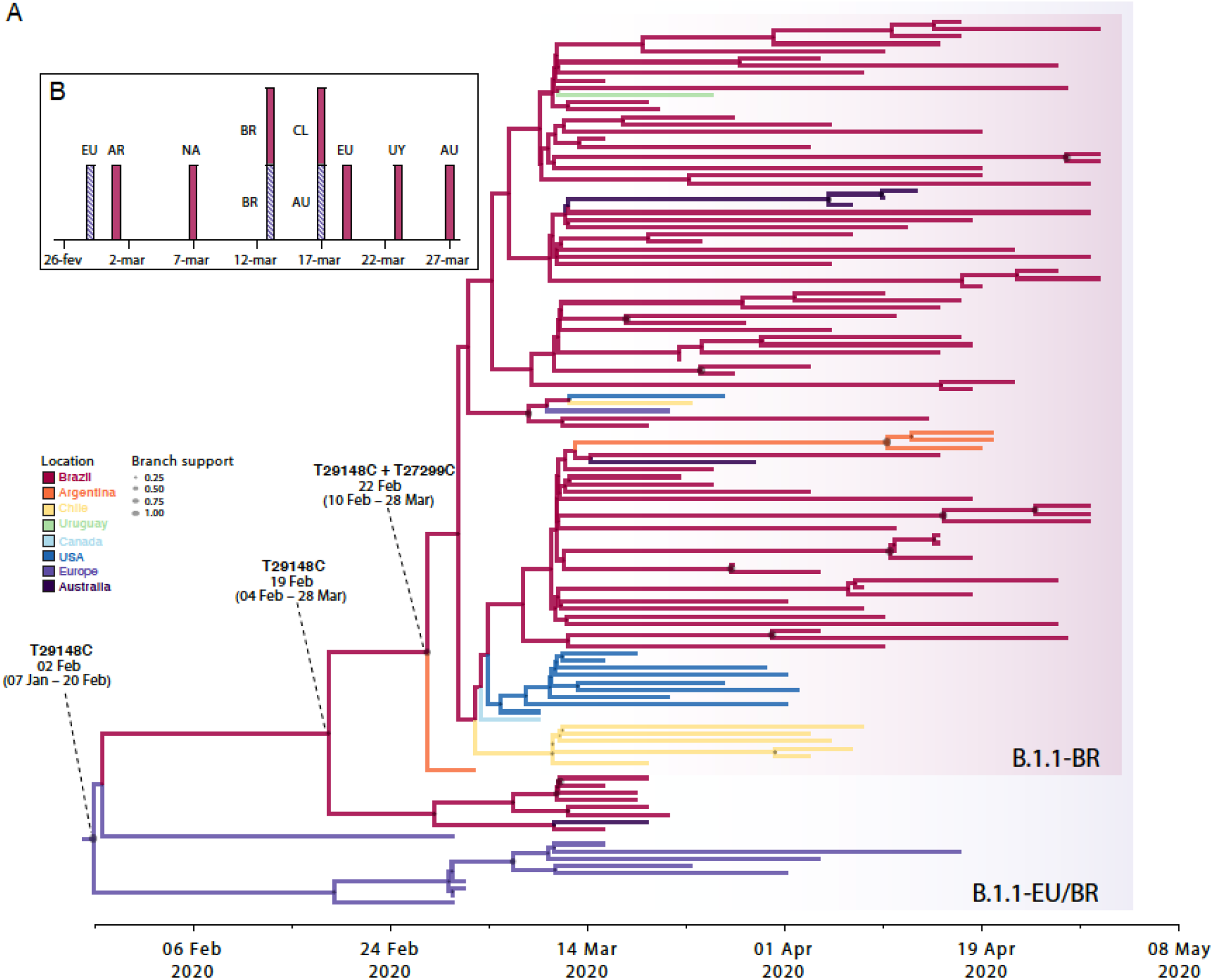
Spatiotemporal dissemination of the SARS-CoV-2 clades B.1.1.EU/BR and B.1.1.BR. A) Time-scaled Bayesian phylogeographic MCC tree of the major B.1.1 lineages circulating in Brazil. Branches are colored according to the most probable location state of their descendent nodes as indicated at the legend. Circles size at internal nodes is proportional to the corresponding posterior probability support as indicated at the legend. The inferred TMRCA (based on the median of the posterior heights) and nucleotide substitutions fixed at ancestral key nodes are shown. Shaded boxes highlight the position of the clades B.1.1.EU/BR and B.1.1.BR. The tree is automatically rooted under the assumption of a strict molecular clock and all horizontal branch lengths are drawn to a scale of years. B) Timeline of the earliest detection of clades B.1.1.EU/BR (blue bars) and B.1.1.BR (red bars) in Europe (EU), North America (NA), Australia (AU), Argentina (AR), Brazil (BR), Chile (CL) and Uruguay (UY).

## Discussion

Our genomic survey identified a major SARS-CoV-2 B.1.1 lineage, here designated as B.1.1.BR, that seems to be responsible for a substantial fraction of the community viral transmissions in Brazil. This lineage harbors two non-synonymous synapomorphic mutations at positions T27299C and T29148C located at the ORF6 (I33T) and the Nucleocapsid protein (I292T), respectively. Basal to this clade, we identified a group of Brazilian and European sequences composing a paraphyletic clade, designated as B.1.1.EU/BR, that only carry the synapomorphic mutation T29148C and seems to represent and evolutionary intermediate between clades B.1.1 and B.1.1.BR.

Our phylogeographic reconstruction supports that clade B.1.1.EU/BR most probably arose in Europe (*PSP* = 0.64) around 2^nd^ February and was introduced into Brazil a couple of weeks later, where it spread and rapidly fixed the T27299C mutation, originating the clade B.1.1.BR (**Fig. 4A**). This evolutionary pattern agrees with the earlier detection of the clade B.1.1.EU/BR in Western Europe (28^th^ February) than in Brazil (13^th^ March) (**Fig. 3B**); but the extremely low prevalence of the clade B.1.1.EU/BR in Europe (<1% of total SARS-CoV-2 sequences) makes this transmission history a highly unlikely epidemiological scenario. Once our phylogeographic analysis also estimated Brazil as a putative ancestral state at the root of clade B.1.1.EU/BR (*PSP* = 0.35), an alternative hypothesis would be that a highly prevalent B.1.1 strain was introduced from Western Europe into Brazil before 2^nd^ February and that synapomorphic mutations T29148C and T27299C were fixed at sequential steps during subsequent virus local spread (**Fig. 4B**). The relative high prevalence of clade B.1.1.EU/BR in some Brazilian locations makes the dissemination of this lineage from Brazil to Western Europe a quite plausible transmission scenario. Retrospective analyses of Brazilian samples obtained from individuals with severe acute respiratory disease during February might provide unique insights to resolve the origin of the clade B.1.1.EU/BR.

**Figure 4.**
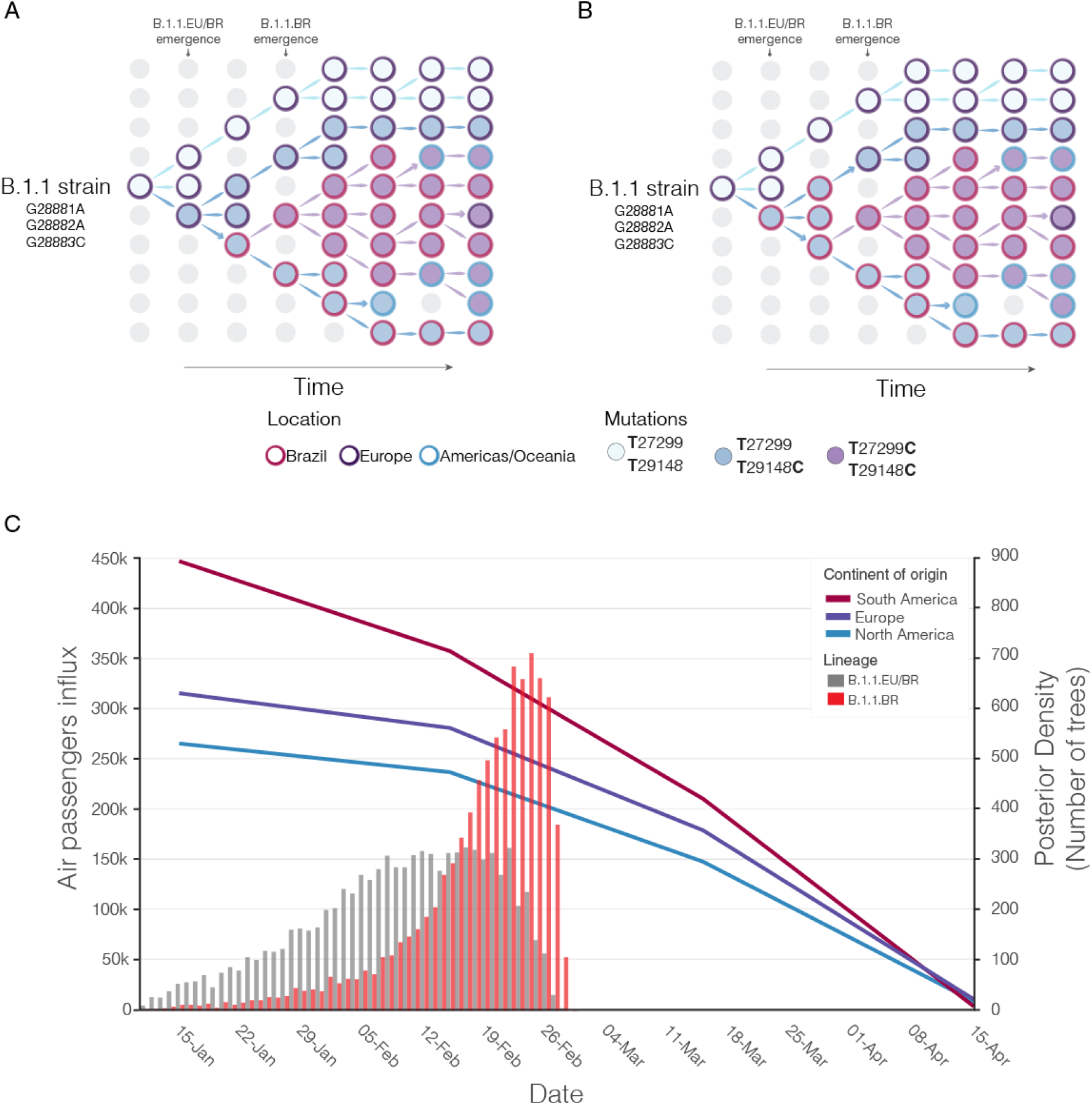
Putative origin and transmission history of the SARS-CoV-2 clades B.1.1.EU/BR and B.1.1.BR. A) Diagrams showing two alternative scenarios for the origin and dissemination of clades B.1.1.EU/BR and B.1.1.BR. The left panel depicts the hypothetical scenario where a B.1.1.EU/BR strain carrying the mutation T29148C was introduced into Brazil from Europe and after a period of local transmission in Brazil arose the B.1.1.BR variant carrying the mutation T27299C, which dispersed all over the country and from Brazil to other countries in the Americas and Oceania. The right panel depicts the hypothetical scenario where a B.1.1 strain was introduced from Europe to Brazil and mutations T29148C and T27299C arose at sequential steps during local transmission. According to this second scenario, Brazil was the epicenter of dissemination of both clades B.1.1.EU/BR and B.1.1.BR to other countries in Europe, the Americas and Oceania. B) Graphic showing the monthly number of international air passengers from South America, North America and Europe that arrived in Brazil during 2020 (available at: https://www.anac.gov.br) (left hand axis) along with probability density of TMRCA estimates for clades B.1.1.EU/BR (gray) and B.1.1.BR (red).

Brazil was traced as the source location of the clade B.1.1.BR with high probability (*PSP* = 0.95) in our phylogeographic reconstruction. The earliest B.1.1.BR sequence currently available, however, is an Argentinean sequence sampled on 1^st^ March 2020; while the earliest detection of the clade B.1.1.BR in Brazil occurred in two samples isolated in the Distrito Federal on 13^th^ March 2020 (**Fig. 3B**). Of note, none of the 30 Brazilian genomes analyzed between 25^th^ February and 12^th^ March, including 12 B.1.1 genomes from imported cases, belong to the clade B.1.1.BR. This suggests that clade B.1.1.BR may have arisen in a Brazilian state that was not included in our dataset and/or that this lineage circulated cryptically for several weeks before being detected in symptomatic carriers. The nearly simultaneous detection of the clade B.1.1.BR in distant states from the Central-Western (Federal District, 13^th^ March), Northern (Amapa, 17^th^ March) and Southeastern (Rio de Janeiro, 20^th^ March) Brazilian regions, supports the second hypothesis and further suggests a wide geographic spread of this Brazilian lineage. Our results also suggest a high geographic compartmentalization of SARS-CoV-2 genetic diversity within Brazil. Considering the three most populated and densely sampled states of Brazil, the clade B.1.1.EU/BR was only detected in Minas Gerais, the clade B.1.1.BR only in Rio the Janeiro, and none of those in Sao Paulo.

Our analyses support that the clade B.1.1.BR not only spread within Brazil but was also exported from Brazil to neighboring South American countries and also to more distant countries (i.e. Canada, USA, UK and Australia). The chance introduction of SARS-CoV-2 strains from Western Europe into Brazil during February and the subsequent exportation of Brazilian SARS-CoV-2 lineages to neighboring South American countries, Western Europe and North America during following weeks agrees with the high influx of tourists from those regions into Brazil during January and February (**Fig. 4C**). Our findings support that when first control measures for international travels were implemented in Brazil around the middle March, the clades B.1.1.EU/BR and B.1.1.BR were already established in the country and also spread from Brazil to other countries. Of note, no B.1.1.EU/BR or B.1.1.BR sequences were detected in Europe, North America or Oceania after 15^th^ April, coinciding with a sharp decrease in the influx of international air travels to Brazil (**Fig. 4C**) that might have greatly reduced the chance of exportation of SARS-CoV-2 Brazilian lineages to other countries.

Our phylogeographic reconstruction suggests that the clade B.1.1.BR might have seeded secondary outbreaks in Argentina, Chile, Australia and the US, but those findings should be interpreted with caution because of the low support of local clusters. Although high-quality full genomes of SARS-CoV-2 currently available contain enough information to allow reliable phylogenetic inferences, the low genetic diversity of within-country (or regional) transmission clusters imposes a serious limitation for accurate phylogeographic reconstructions ^39,40^. Indeed, the MC test supports a random phylogenetic clustering of B.1.1.EU/BR and B.1.1.BR strains from most locations, with exception of Brazil, Argentina and Europe (**Supplementary Table 4**). The B.1.1.BR sequences sampled at different Brazilian states were also highly similar or identical, making it difficult to trace with precision the origin and within-country fluxes of this viral clade during the early epidemic phase in Brazil. Another important limitation of our study is the uneven spatial and temporal sampling scheme. Most SARS-CoV-2 sequences recovered in the present study were from the Rio de Janeiro state and might thus not represent the viral diversity in other Brazilian states. More accurate reconstructions of the origin and regional spread of the clade B.1.1.BR will require a denser sampling from Brazil and neighboring South American countries, particularly during the very early phase of the epidemic.

In summary, this study reveals the existence of a major SARS-CoV-2 B.1.1 lineage associated with community transmission in Brazil and widespread in a national scale. This major B.1.1 Brazilian lineage emerged in Brazil in February 2020, probably before the detection of the first imported SARS-CoV-2 case in the country, and reached different Brazilian regions by the middle of March 2020. Continuous efforts for widespread sequencing of SARS-CoV-2 may provide unique insight about its local dissemination in Brazil and other South American countries.

## Acknowledgements

The authors wish to thank all the health care workers and scientists, who have worked hard to deal with this pandemic threat, the GISAID team and all the submitters of the database. GISAID acknowledgment tables containing sequences used in this study are in **Supplementary Tables 5** (South American SARS-CoV-2 genomes) and **Supplementary Table 6** (Global SARS-CoV 2 genomes B.1.1 lineage). Locally, we acknowledge the Respiratory Viruses Genomic Surveillance Network of the General Laboratory Coordination (CGLab) of the Brazilian Ministry of Health (MoH), Brazilian Central Laboratory States (LACENs), and local surveillance teams for the partnership in the viral surveillance in Brazil.

## Funding support

CGLab/MoH (General Laboratories Coordination of Brazilian Ministry of Health) and CVSLR/FIOCRUZ (Coordination of Health Surveillance and Reference Laboratories of Oswaldo Cruz Foundation).

## Supplementary Material

**Supplementary Figure 1.**
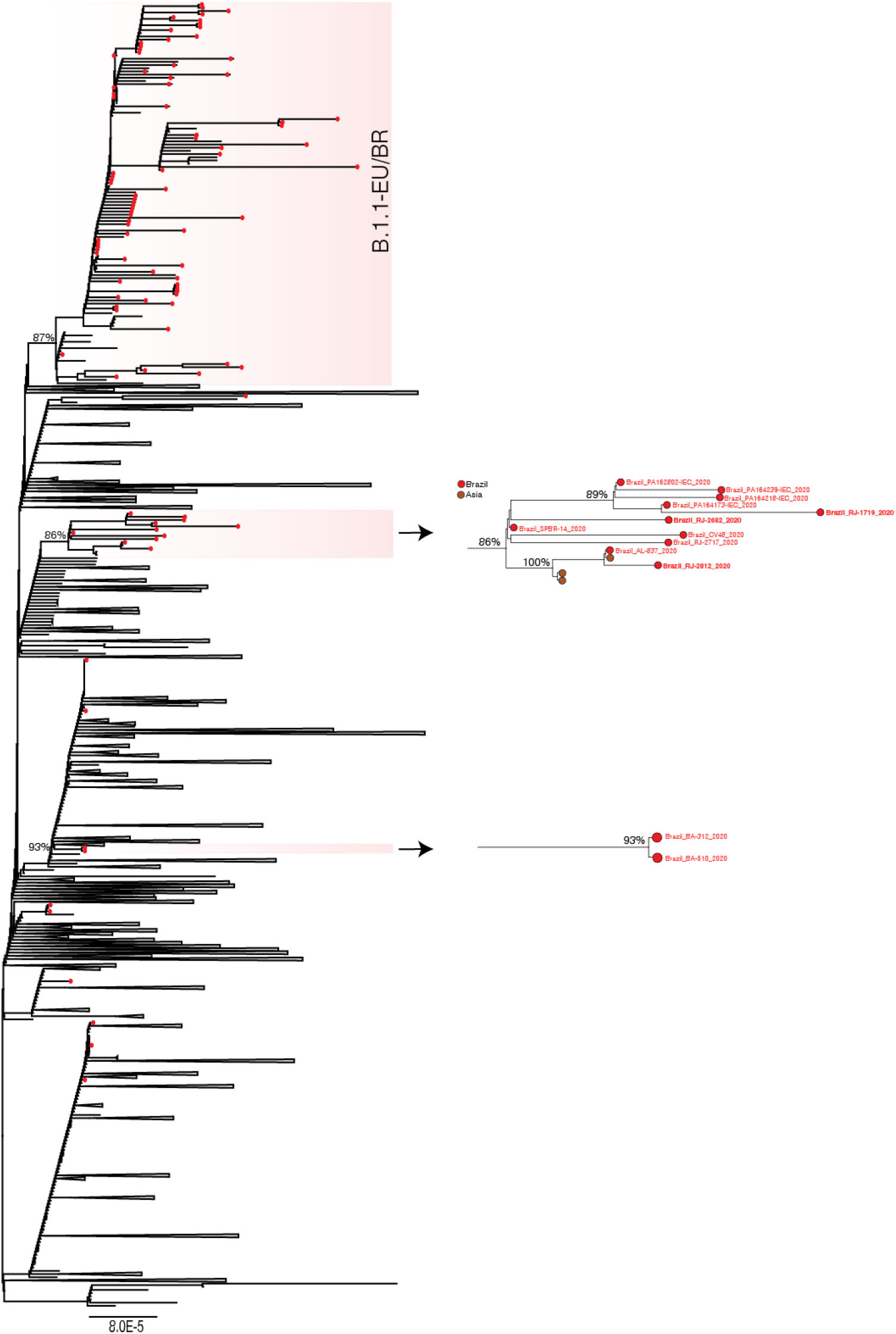
Phylogenetic relationships of SARS-CoV-2 B.1.1 Brazilian and global strains. ML phylogenetic tree of 87 B.1.1 Brazilian genomes obtained in this survey (red circles) along with 3,764 B.1.1 worldwide reference sequences from GISAID database. Shaded box highlights the position of major Brazilian clusters B.1.1.EU/BR and B.1.1.BR, and a close view of each cluster is showed. Names of Brazilian SARS-CoV-2 genomes generated in this study (red tips) are shown. Only node supports (SH-*aLRT*) above 70% are shown. Tree was rooted on midpoint and branch lengths are drawn to scale with the bars at the bottom indicating nucleotide substitutions per site.

**Supplementary Figure 2.**
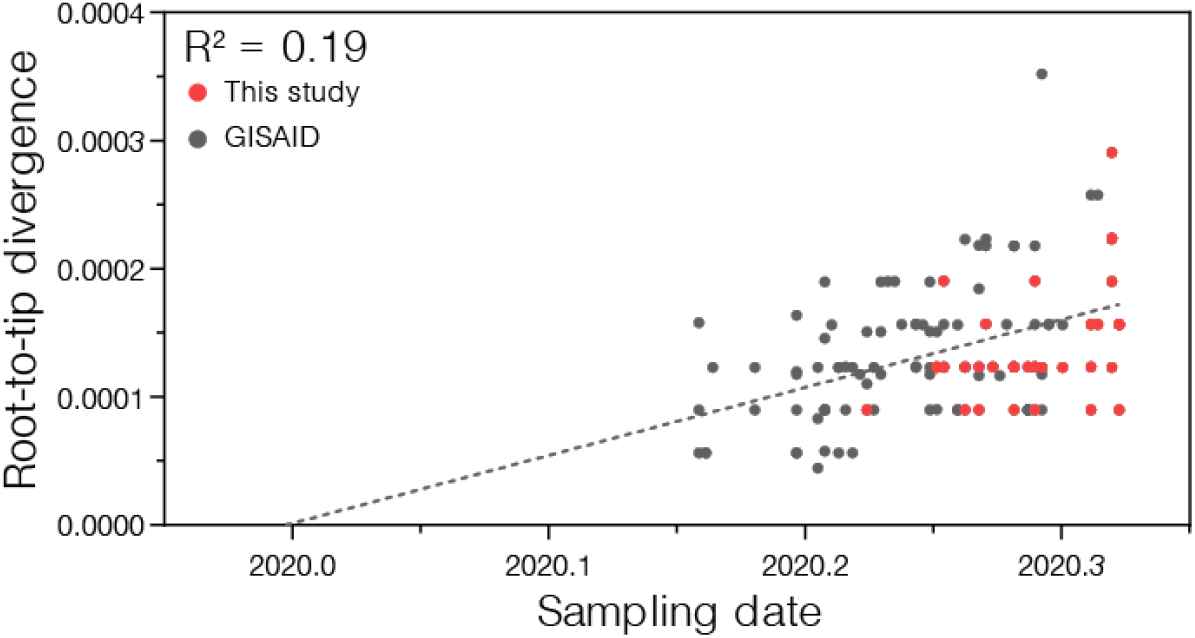
Linear regression analysis between the sampling date of each viral sequence and the root-to-tip divergence (genetic distance of that sequence to the tree root) of a ML phylogeny of the SARS-CoV-2 clades B.1.1.EU/BR and B.1.1.BR. SARS-CoV-2 genomes generated in this study (red) were combined with other genomes available on the GISAID database (gray).

**Supplementary Table 1.** Clinical and epidemiological data associated with SARS-CoV-2 genomes obtained in this study.

| Virus name | Accession number<br>GISAID | Clinical<br>Specimen | Pangolin<br>lineage* | Town | Collection<br>date | Onset<br>symptoms | Travel<br>history | Gender | Age | Clinical<br>outcome |
| --- | --- | --- | --- | --- | --- | --- | --- | --- | --- | --- |
| hCoV-19/Brazil/ES-225/2020 | EPI_ISL_415128 | NPS | B.2.1 | Vila Velha | 29/02/2020 | 22/02/2020 | Italy | F | 37y | inpatient |
| hCoV-19/Brazil/BA-312/2020 | EPI_ISL_415105 | NPS | B.1.1 | Feira de Santana | 04/03/2020 | 26/02/2020 | Italy | F | 34y | inpatient |
| hCoV-19/Brazil/RJ-314/2020 | EPI_ISL_414045 | NPS | B.1 | Rio de Janeiro | 04/03/2020 | 02/03/2020 | Italy | F | 52y | outpatient |
| hCoV-19/Brazil/RJ-352/2020 | EPI_ISL_427299 | NPS | A.2 | Niteroi | 05/03/2020 | unknown | unknown | M | 27y | unknown |
| hCoV-19/Brazil/RJ-477/2020 | EPI_ISL_427300 | NPS | B.1.1 | Rio de Janeiro | 11/03/2020 | 10/03/2020 | Europe | M | 48y | outpatient |
| hCoV-19/Brazil/RJ-477i/2020 | EPI_ISL_427301 | Vero E6 | B.1.1 | Rio de Janeiro | 11/03/2020 | 10/03/2020 | Europe | M | 48y | outpatient |
| hCoV-19/Brazil/BA-510/2020 | EPI_ISL_427293 | NPS | B.1.1 | Feira de Santana | 06/03/2020 | 03/03/2020 | unknown | F | 41y | unknown |
| hCoV-19/Brazil/DF-615i/2020 | EPI_ISL_427294 | Vero E6 | B.1.1.BR | Brasilia | 13/03/2020 | unknown | unknown | M | unknown | unknown |
| hCoV-19/Brazil/DF-619i/2020 | EPI_ISL_427295 | Vero E6 | B.1.1.BR | Brasilia | 13/03/2020 | unknown | unknown | M | unknown | unknown |
| hCoV-19/Brazil/RJ-763/2020 | EPI_ISL_427302 | NPS | B.1.1.BR | Petropolis | 20/03/2020 | 19/03/2020 | no | F | 47y | outpatient |
| hCoV-19/Brazil/SC-766/2020 | EPI_ISL_427305 | NPS | B.6 | Joinville | 10/03/2020 | 08/03/2020 | Europe and Africa | M | 57y | unknown |
| hCoV-19/Brazil/SC-769/2020 | EPI_ISL_427306 | NPS | B.1.1 | Florianopolis | 10/03/2020 | 04/02/2020 | Europe | F | 59y | outpatient |
| hCoV-19/Brazil/RJ-818/2020 | EPI_ISL_427303 | NPS | B.1.1.BR | Rio de Janeiro | 25/03/2020 | 22/03/2020 | no | F | 54y | outpatient |
| hCoV-19/Brazil/AL-837/2020 | EPI_ISL_427292 | NPS | B.1.1 | Maceio | 18/03/2020 | 15/03/2020 | unknown | M | 65y | unknown |
| hCoV-19/Brazil/DF-861/2020 | EPI_ISL_427296 | NPS | B.1.1.BR | Brasilia | 23/03/2020 | unknown | unknown | M | 55y | unknown |
| hCoV-19/Brazil/DF-862/2020 | EPI_ISL_427297 | NPS | B.1.1.BR | Brasilia | 23/03/2020 | unknown | unknown | F | 25y | unknown |
| hCoV-19/Brazil/RJ-872/2020 | EPI_ISL_427304 | NPS | B.1.1 | Rio de Janeiro | 26/03/2020 | unknown | unknown | F | 60y | unknown |
| hCoV-19/Brazil/DF-891/2020 | EPI_ISL_427298 | NPS | B.1.1.BR | Brasilia | 22/03/2020 | 16/03/2020 | unknown | F | 61y | Deceased |
| hCoV-19/Brazil/RJ-1056/2020 | EPI_ISL_456072 | NPS | B.1.1.BR | Rio de Janeiro | 01/04/2020 | 30/03/2020 | no | F | 31y | outpatient |
| hCoV-19/Brazil/RJ-1058/2020 | EPI_ISL_456073 | NPS | B.1.1.BR | Rio de Janeiro | 01/04/2020 | 25/03/2020 | no | M | 27y | outpatient |
| hCoV-19/Brazil/RJ-1065/2020 | EPI_ISL_456074 | NPS | B.1.1.BR | Rio de Janeiro | 01/04/2020 | asymptomatic | no | F | 85y | outpatient |
| hCoV-19/Brazil/RJ-1402/2020 | EPI_ISL_456079 | NPS | B.1.1.BR | Rio de Janeiro | 03/04/2020 | 30/03/2020 | no | F | 55y | outpatient |
| hCoV-19/Brazil/RJ-1464/2020 | EPI_ISL_456080 | NPS | B.1.1.BR | Petropolis | 06/04/2020 | unknown | no | M | 30y | outpatient |
| hCoV-19/Brazil/RJ-1466/2020 | EPI_ISL_456081 | NPS | B.1.1.BR | Rio de Janeiro | 06/04/2020 | 03/04/2020 | no | F | 54y | outpatient |
| hCoV-19/Brazil/RJ-1701/2020 | EPI_ISL_456086 | NPS | B.1.1.BR | Rio de Janeiro | 08/04/2020 | 05/04/2020 | no | M | 60y | outpatient |
| hCoV-19/Brazil/RJ-1702/2020 | EPI_ISL_456087 | NPS | B.1.1.BR | Rio de Janeiro | 08/04/2020 | 04/04/2020 | no | M | 40y | outpatient |
| hCoV-19/Brazil/RJ-1902/2020 | EPI_ISL_456090 | NPS | B.1.1.BR | Rio de Janeiro | 09/04/2020 | 06/04/2020 | no | F | 39y | outpatient |
| hCoV-19/Brazil/RJ-1921/2020 | EPI_ISL_456091 | NPS | B.1.1.BR | Rio de Janeiro | 09/04/2020 | 06/04/2020 | no | F | 70y | outpatient |
| hCoV-19/Brazil/RJ-1923/2020 | EPI_ISL_456092 | NPS | B.1.1.BR | Rio de Janeiro | 10/04/2020 | 08/04/2020 | no | M | 5m | unknown |
| hCoV-19/Brazil/RJ-1927/2020 | EPI_ISL_456093 | NPS | B.1.1.BR | Rio de Janeiro | 12/04/2020 | 10/04/2020 | no | M | 46y | outpatient |
| hCoV-19/Brazil/RJ-1948/2020 | EPI_ISL_456095 | NPS | B.1.1.BR | Rio de Janeiro | 13/04/2020 | 12/04/2020 | no | F | 45y | inpatient |
| hCoV-19/Brazil/RJ-1952/2020 | EPI_ISL_456096 | NPS | B.1.1.BR | Rio de Janeiro | 13/04/2020 | 11/04/2020 | no | M | 29y | outpatient |
| hCoV-19/Brazil/RJ-2057/2020 | EPI_ISL_456102 | NPS | B.1.1.BR | Rio de Janeiro | 16/04/2020 | 11/04/2020 | no | M | 29y | outpatient |
| hCoV-19/Brazil/RJ-899/2020 | EPI_ISL_456071 | NPS | B.1.1 | Rio de Janeiro | 30/03/2020 | 24/03/2020 | no | M | 42y | outpatient |
| hCoV-19/Brazil/RJ-1100/2020 | EPI_ISL_456075 | NPS | B.1.1.BR | Belford Roxo | 02/04/2020 | 30/01/2020 | unknown | M | 30y | outpatient |
| hCoV-19/Brazil/RJ-1111/2020 | EPI_ISL_456076 | NPS | B.1.1.BR | Rio de Janeiro | 25/03/2020 | 22/03/2020 | unknown | M | 59y | unknown |
| hCoV-19/Brazil/RJ-1119/2020 | EPI_ISL_456077 | NPS | B.1.1.BR | Rio de Janeiro | 25/05/2020 | 20/03/2020 | unknown | M | 55y | unknown |
| hCoV-19/Brazil/RJ-1555/2020 | EPI_ISL_467344 | NPS | B.2.2 | Petropolis | 02/04/2020 | 27/03/2020 | unknown | M | 31y | unknown |
| hCoV-19/Brazil/RJ-1574/2020 | EPI_ISL_467345 | NPS | B.1.1.BR | Rio de Janeiro | 02/04/2020 | 25/03/2020 | unknown | M | 82y | unknown |
| hCoV-19/Brazil/RJ-1595/2020 | EPI_ISL_467346 | NPS | B.2.2 | Rio de Janeiro | 02/04/2020 | 28/03/2020 | unknown | F | 83y | unknown |
| hCoV-19/Brazil/RJ-1600/2020 | EPI_ISL_456082 | NPS | B.1.1.BR | Teresopolis | 02/04/2020 | 24/03/2020 | unknown | M | 68y | unknown |
| hCoV-19/Brazil/RJ-1627/2020 | EPI_ISL_456083 | NPS | B.1.1.BR | Nova Iguaçu | 03/04/2020 | 30/03/2020 | unknown | F | 44y | unknown |
| hCoV-19/Brazil/RJ-1690/2020 | EPI_ISL_456084 | NPS | B.1.1.BR | Rio de Janeiro | 08/04/2020 | 07/04/2020 | unknown | F | 42y | outpatient |
| hCoV-19/Brazil/RJ-1691/2020 | EPI_ISL_456085 | NPS | B.1.1.BR | Rio de Janeiro | 08/04/2020 | 06/04/2020 | unknown | M | 42y | outpatient |
| hCoV-19/Brazil/RJ-1719/2020 | EPI_ISL_456088 | NPS | B.1.1 | Itaboraí | 06/04/2020 | 05/04/2020 | unknown | F | 36y | unknown |
| hCoV-19/Brazil/RJ-1901/2020 | EPI_ISL_456089 | NPS | B.1.1.BR | Rio de Janeiro | 09/04/2020 | 06/04/2020 | unknown | F | 59y | outpatient |
| hCoV-19/Brazil/RJ-1943/2020 | EPI_ISL_456094 | NPS | B.1.1.BR | Rio de Janeiro | 13/04/2020 | 11/04/2020 | unknown | F | 38y | outpatient |
| hCoV-19/Brazil/RJ-1966/2020 | EPI_ISL_456097 | NPS | B.1.1.BR | Rio de Janeiro | 13/04/2020 | unknown | unknown | M | unknown | outpatient |
| hCoV-19/Brazil/RJ-2000/2020 | EPI_ISL_456098 | NPS | B.1.1.BR | Rio de Janeiro | 13/04/2020 | 13/04/2020 | unknown | F | 29y | unknown |
| hCoV-19/Brazil/RJ-2007/2020 | EPI_ISL_456099 | NPS | B.1.1.BR | Rio de Janeiro | 13/04/2020 | 10/04/2020 | unknown | F | 58y | unknown |
| hCoV-19/Brazil/RJ-2033/2020 | EPI_ISL_456100 | NPS | B.1.1.BR | Rio de Janeiro | 15/04/2020 | 09/04/2020 | unknown | F | 37y | unknown |
| hCoV-19/Brazil/RJ-2044/2020 | EPI_ISL_456101 | NPS | B.1.1.BR | Duque de Caxias | 15/04/2020 | 14/04/2020 | unknown | F | 28y | unknown |
| hCoV-19/Brazil/RJ-2062/2020 | EPI_ISL_456103 | NPS | B.1.1 | Rio de Janeiro | 16/04/2020 | 13/04/2020 | unknown | F | 49y | unknown |
| hCoV-19/Brazil/RJ-2072/2020 | EPI_ISL_456104 | NPS | B.1.1.BR | Rio de Janeiro | 16/04/2020 | 12/04/2020 | unknown | F | 51y | unknown |
| hCoV-19/Brazil/RJ-2077/2020 | EPI_ISL_456105 | NPS | B.1.1.BR | Rio de Janeiro | 16/04/2020 | unknown | unknown | F | 38y | unknown |
| hCoV-19/Brazil/RJ-2078/2020 | EPI_ISL_456106 | NPS | B.1.1.BR | Rio de Janeiro | 16/04/2020 | 14/04/2020 | unknown | F | 34y | unknown |
| hCoV-19/Brazil/RJ-2091/2020 | EPI_ISL_467347 | NPS | B.1.1.BR | Rio de Janeiro | 16/04/2020 | 12/04/2020 | unknown | F | 51y | unknown |
| hCoV-19/Brazil/RJ-2195/2020 | EPI_ISL_467348 | NPS | B.1.1.BR | Rio de Janeiro | 17/04/2020 | 14/04/2020 | unknown | M | 45y | unknown |
| hCoV-19/Brazil/RJ-2197/2020 | EPI_ISL_467349 | NPS | B.1.1.BR | Rio de Janeiro | 17/04/2020 | 14/04/2020 | unknown | M | 32y | unknown |
| hCoV-19/Brazil/RJ-2208/2020 | EPI_ISL_467350 | NPS | B.1.1.BR | Rio de Janeiro | 17/04/2020 | 15/04/2020 | unknown | M | 43y | unknown |
| hCoV-19/Brazil/RJ-2233/2020 | EPI_ISL_467351 | NPS | B.1.1.BR | Rio de Janeiro | 17/04/2020 | 13/04/2020 | unknown | F | 30y | unknown |
| hCoV-19/Brazil/RJ-2422/2020 | EPI_ISL_467352 | NPS | B.1.1.BR | Queimados | 20/04/2020 | 18/04/2020 | unknown | F | 27y | inpatient |
| hCoV-19/Brazil/RJ-2669/2020 | EPI_ISL_467353 | NPS | B.1.1.BR | Niteroi | 24/04/2020 | 21/04/2020 | unknown | F | 25y | outpatient |
| hCoV-19/Brazil/RJ-2676/2020 | EPI_ISL_467354 | NPS | B.1.5 | Rio de Janeiro | 24/04/2020 | 23/04/2020 | unknown | M | 48y | unknown |
| hCoV-19/Brazil/RJ-2678/2020 | EPI_ISL_467355 | NPS | B.1.1.BR | Rio de Janeiro | 24/04/2020 | 23/04/2020 | unknown | M | 29y | unknown |
| hCoV-19/Brazil/RJ-2682/2020 | EPI_ISL_467356 | NPS | B.1.1 | Rio de Janeiro | 24/04/2020 | 22/04/2020 | unknown | F | 29y | unknown |
| hCoV-19/Brazil/RJ-2683/2020 | EPI_ISL_467357 | NPS | B.1.1.BR | Rio de Janeiro | 24/04/2020 | 20/04/2020 | unknown | F | 31y | unknown |
| hCoV-19/Brazil/RJ-2696/2020 | EPI_ISL_467358 | NPS | B.1.1.BR | Rio de Janeiro | 24/04/2020 | 19/04/2020 | unknown | F | 65y | unknown |
| hCoV-19/Brazil/RJ-2717/2020 | EPI_ISL_467359 | NPS | B.1.1 | Rio de Janeiro | 24/04/2020 | 21/04/2020 | unknown | F | 36y | unknown |
| hCoV-19/Brazil/RJ-2733/2020 | EPI_ISL_467360 | NPS | B.1.1.BR | Duque de Caxias | 25/04/2020 | 22/04/2020 | unknown | F | 49y | unknown |
| hCoV-19/Brazil/RJ-2769/2020 | EPI_ISL_467361 | NPS | B.1.1.BR | Rio de Janeiro | 27/04/2020 | 22/04/2020 | unknown | F | 30y | unknown |
| hCoV-19/Brazil/RJ-2770/2020 | EPI_ISL_467362 | NPS | B.1.1.BR | Rio de Janeiro | 27/04/2020 | 22/04/2020 | unknown | F | 30y | unknown |
| hCoV-19/Brazil/RJ-2776/2020 | EPI_ISL_467363 | NPS | B.1.1.BR | Rio de Janeiro | 27/04/2020 | 21/04/2020 | unknown | M | 64y | unknown |
| hCoV-19/Brazil/RJ-2777/2020 | EPI_ISL_467364 | NPS | B.1.1.BR | Nova Iguaçu | 25/04/2020 | 25/04/2020 | unknown | F | 34y | unknown |
| hCoV-19/Brazil/RJ-2811/2020 | EPI_ISL_467365 | NPS | B.1.1.BR | Rio de Janeiro | 27/04/2020 | 25/04/2020 | unknown | M | 44y | unknown |
| hCoV-19/Brazil/RJ-2812/2020 | EPI_ISL_467366 | NPS | B.1.1 | Rio de Janeiro | 27/04/2020 | 20/04/2020 | unknown | M | 38y | unknown |
| hCoV-19/Brazil/RJ-2822/2020 | EPI_ISL_467367 | NPS | B.1.1.BR | Niteroi | 27/04/2020 | 23/04/2020 | unknown | M | 62y | inpatient |
| hCoV-19/Brazil/RJ-2840/2020 | EPI_ISL_467368 | NPS | B.1.1.BR | Rio de Janeiro | 28/04/2020 | 25/04/2020 | unknown | M | 36y | unknown |
| hCoV-19/Brazil/RJ-2844/2020 | EPI_ISL_467369 | NPS | B.1.1.BR | Rio de Janeiro | 28/04/2020 | 24/04/2020 | unknown | F | 29y | outpatient |
| hCoV-19/Brazil/RJ-2847/2020 | EPI_ISL_467370 | NPS | B.1.1.BR | Rio de Janeiro | 28/04/2020 | 22/04/2020 | unknown | F | 35y | unknown |
| hCoV-19/Brazil/RJ-2868/2020 | EPI_ISL_467371 | NPS | B.1.1.BR | Rio de Janeiro | 28/04/2020 | 25/04/2020 | unknown | F | 49y | unknown |
| hCoV-19/Brazil/AP-161167-IEC/2020 | EPI_ISL_450873 | Sputum | B.1.1.BR | Macapa | 17/03/2020 | 12/03/2020 | No | F | 36y | unknown |
| hCoV-19/Brazil/PA-161548-IEC/2020 | EPI_ISL_450874 | NPS | B.1.1.BR | Maraba | 20/03/2020 | 16/03/2020 | Sao Paulo | F | 28y | unknown |
| hCoV-19/Brazil/AP-162741-IEC/2020 | EPI_ISL_458138 | Sputum | B.1.1.BR | Macapa | 03/04/2020 | 02/04/2020 | unknown | F | 37y | unknown |
| hCoV-19/Brazil/AC-162535-IEC/2020 | EPI_ISL_458139 | NPS | B.1.1.BR | Rio Branco | 18/03/2020 | 15/03/2020 | No | M | 81y | unknown |
| hCoV-19/Brazil/PA-162802-IEC/2020 | EPI_ISL_458140 | NPS | B.1.1 | Belem | 07/04/2020 | 06/04/2020 | No | M | 63y | inpatient |
| hCoV-19/Brazil/PA-164239-IEC/2020 | EPI_ISL_458141 | NPS | B.1.1 | Ananindeua | 26/04/2020 | 24/04/2020 | No | M | 38y | outpatient |
| hCoV-19/Brazil/AP-162966-IEC/2020 | EPI_ISL_458142 | NPS | B.1.1.BR | Macapa | 05/04/2020 | 05/04/2020 | No | F | 37y | unknown |
| hCoV-19/Brazil/AP-164082-IEC/2020 | EPI_ISL_458143 | Sputum | B.1.1.BR | Macapa | 15/04/2020 | 13/04/2020 | unknown | M | 44y | unknown |
| hCoV-19/Brazil/AP-163972-IEC/2020 | EPI_ISL_458144 | Sputum | B.1.1.BR | Macapa | 15/04/2020 | 12/04/2020 | unknown | M | 44y | unknown |
| hCoV-19/Brazil/AP-164346-IEC/2020 | EPI_ISL_458145 | Sputum | B.1.1.BR | Macapa | 20/04/2020 | 28/04/2020 | unknown | F | 62y | unknown |
| hCoV-19/Brazil/PA-164173-IEC/2020 | EPI_ISL_458146 | NPS | B.1.1 | Ananindeua | 23/04/2020 | 20/04/2020 | unknown | F | 38y | unknown |
| hCoV-19/Brazil/PA-164218-IEC/2020 | EPI_ISL_458147 | NPS | B.1.1 | Belem | 24/04/2020 | unknown | unknown | M | 45y | unknown |
| hCoV-19/Brazil/PA-164684-IEC/2020 | EPI_ISL_458148 | NPS | B.1.1.BR | Belem | 27/04/2020 | 26/04/2020 | no | M | 38y | unknown |
| hCoV-19/Brazil/MA-163069-IEC/2020 | EPI_ISL_458149 | NPS | B.1.1.BR | Sao Luis | 06/04/2020 | unknown | unknown | F | 40y | unknown |

NPS, Nasopharyngeal swab

**Supplementary Table 2.** Prevalence of SARS-CoV-2 lineages B.1.1.EU/BR and B.1.1.BR across countries.

| Region | Country | Total SARS-Cov-2 | Lineage B.1.1.EU/BR | Lineage B.1.1.BR |
| --- | --- | --- | --- | --- |
| South America | Brazil | 170 | 10 (6%) | 74 (44%) |
|  | Argentina | 29 | - | 4 (14%) |
|  | Chile | 153 | - | 7 (5%) |
|  | Uruguay | 45 | - | 1 (2%) |
| North America | Canada | 227 | - | 1 (<1%) |
|  | US | 7,605 | - | 10 (<1%) |
| Oceania | Australia | 1,899 | 1 (<1%) | 4 (<1%) |
| Europe | United Kingdom | 18,391 | 4 (<1%) | 1 (<1%) |
|  | Switzerland | 325 | 4 (1%) | - |
|  | Netherlands | 840 | 2 (<1%) | - |

**Supplementary Table 3.** Prevalence of SARS-CoV-2 lineages B.1.1.EU/BR and B.1.1.BR across Brazilian states.

| State | Total SARS-Cov-2 | Lineage B.1.1.EU/BR | Lineage B.1.1.BR |
| --- | --- | --- | --- |
| Rio de Janeiro | 77 | - | 59 (76%) |
| Minas Gerais | 45 | 9 (20%)* | - |
| Sao Paulo | 18 | - | - |
| Distrito Federal | 7 | 1 (14%) | 5 (71%) |
| Amapa | 6 | - | 6 (100%) |
| Para | 6 | - | 2 (33%) |
| Others | 11 | - | 2 (18%) |
| **Total** | **170** | **10 (6%)** | **74 (44%)** |

\* Sequences CV34 and CV45 harbor the substitution T29148C, but displayed an ambiguous nucleotide at position 27299, hence assigned to lineage B.1.1.EU/BR.

**Supplementary Table 4.**
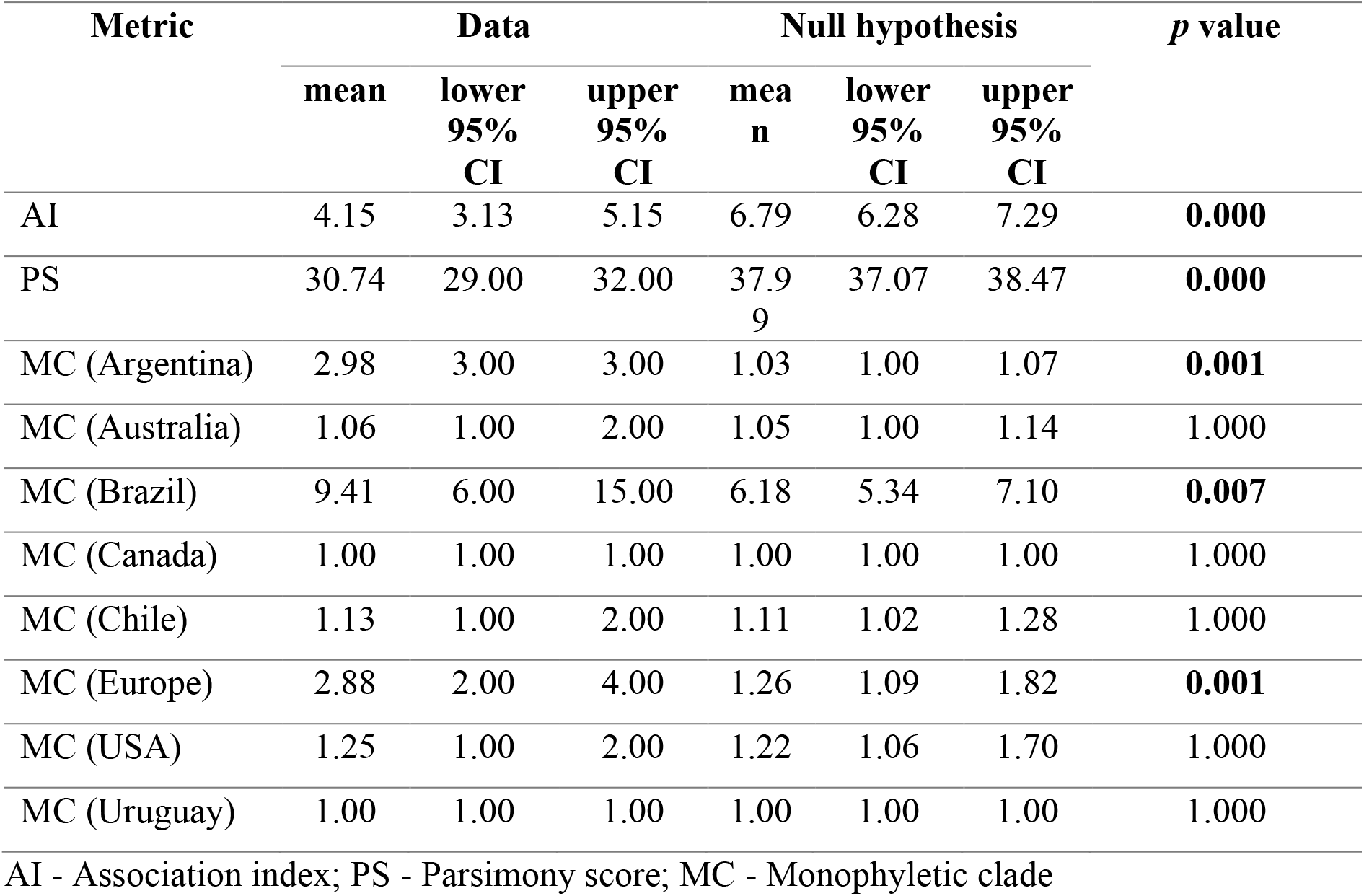
Phylogeny-trait association tests to assess phylogeographic structure of the dataset

| Metric | Data |  |  | Null hypothesis |  |  | *p* value |
| --- | --- | --- | --- | --- | --- | --- | --- |
|  | mean | lower 95% CI | upper 95% CI | mean | lower 95% CI | upper 95% CI |  |
| AI | 4.15 | 3.13 | 5.15 | 6.79 | 6.28 | 7.29 | **0.000** |
| PS | 30.74 | 29.00 | 32.00 | 37.99 | 37.07 | 38.47 | **0.000** |
| MC (Argentina) | 2.98 | 3.00 | 3.00 | 1.03 | 1.00 | 1.07 | **0.001** |
| MC (Australia) | 1.06 | 1.00 | 2.00 | 1.05 | 1.00 | 1.14 | 1.000 |
| MC (Brazil) | 9.41 | 6.00 | 15.00 | 6.18 | 5.34 | 7.10 | **0.007** |
| MC (Canada) | 1.00 | 1.00 | 1.00 | 1.00 | 1.00 | 1.00 | 1.000 |
| MC (Chile) | 1.13 | 1.00 | 2.00 | 1.11 | 1.02 | 1.28 | 1.000 |
| MC (Europe) | 2.88 | 2.00 | 4.00 | 1.26 | 1.09 | 1.82 | **0.001** |
| MC (USA) | 1.25 | 1.00 | 2.00 | 1.22 | 1.06 | 1.70 | 1.000 |
| MC (Uruguay) | 1.00 | 1.00 | 1.00 | 1.00 | 1.00 | 1.00 | 1.000 |

AI - Association index; PS - Parsimony score; MC - Monophyletic clade

**Supplementary Table 5.** GISAID acknowledgment table of South America SARS-CoV-2 genomes.

**Supplementary Table 6.** GISAID acknowledgment table of Global SARS-CoV-2 genomes B.1.1 lineage.

## Notes

### Competing Interest Statement

The authors have declared no competing interest.

